# Inducible transposon mutagenesis for genome-scale forward genetics

**DOI:** 10.1101/2024.05.21.595064

**Authors:** David W. Basta, Ian W. Campbell, Emily J. Sullivan, Julia A. Hotinger, Karthik Hullahalli, Matthew K. Waldor

## Abstract

Transposon insertion sequencing (Tn-seq) is a powerful method for genome-scale functional genetics in bacteria. However, its effectiveness is often limited by a lack of mutant diversity, caused by either inefficient transposon delivery or stochastic loss of mutants due to population bottlenecks. Here, we introduce “InducTn-seq”, which leverages inducible mutagenesis for temporal control of transposition. InducTn-seq generates millions of transposon mutants from a single colony, enabling the sensitive detection of subtle fitness defects and transforming binary classifications of gene essentiality into a quantitative fitness measurement across both essential and non-essential genes. Using a mouse model of infectious colitis, we show that InducTn-seq bypasses a highly restrictive host bottleneck to generate a diverse transposon mutant population from the few cells that initiate infection, revealing the role of oxygen-related metabolic plasticity in pathogenesis. Overall, InducTn-seq overcomes the limitations of traditional Tn-seq, unlocking new possibilities for genome-scale forward genetic screens in bacteria.

## Main

Since its inception in 2009, transposon insertion sequencing (Tn-seq) has been the most widely used method for conducting genome-scale forward genetic screens in bacteria^1–7^. The method involves mutagenesis of a bacterial strain with a randomly integrating transposon followed by quantification of transposon insertion sites across the genome using high-throughput sequencing. A relative paucity of transposon insertions within a locus suggests that the locus is important for bacterial fitness in a specific condition. Tn-seq has been applied in various contexts, including the identification of genes required for bacterial fitness within animal tissues^6–8^.

Tn-seq requires the generation of a diverse transposon mutant population. However, achieving sufficient diversity is often constrained by the efficiency of transposon delivery. The creation of diverse transposon libraries, which typically contain ∼10^5^ unique mutants, can be a labor-intensive, rate-limiting step in conducting Tn-seq screens. Achieving high diversity libraries is especially difficult when working with organisms that are inefficient at taking up exogenous DNA.

In traditional Tn-seq, transposon delivery is usually accompanied by the loss of the transposase during the enrichment of the mutant population, thereby halting further mutagenesis. Consequently, the diversity of the mutant population is not temporally controllable and can be eradicated by severe population bottlenecks, which cause loss of mutants from the population due to random chance, rather than due to the selection of interest. Traditional Tn-seq cannot overcome this “bottleneck problem” since it does not enable mutant generation post-bottleneck. This limitation has curtailed the application of Tn-seq for the discovery of genes that promote pathogen fitness in most animal models because host-imposed bottlenecks often markedly reduce the number of unique mutant cells contributing to the bacterial population within the animal^9–32^. In fact, fewer than 10^3^ bacterial cells, a number insufficient for conducting a genome-scale genetic screen, initiate infection in the majority of animal colonization models^9,12,16,20,22,24–32^.

We hypothesized that temporal control of transposition could overcome limitations to traditional Tn-seq arising from inefficient transposon delivery or loss of library diversity due to host bottlenecks (Fig. 1). Inspired by work using inducible transposase expression^33–36^, we developed a simplified and streamlined approach for generating maximally diverse and temporally controllable mutant populations, which we term “InducTn-seq” because it combines inducible mutagenesis (induction) with transposon insertion sequencing (Tn-seq). We show that InducTn-seq offers several advantages over traditional Tn-seq. These advantages include the ease of generating highly diverse mutant libraries, the sensitive detection of mutants with subtle fitness defects, and the ability to perform quantitative analysis of mutant fitness for genes previously characterized as essential, thereby transforming binary classifications of essentiality. Finally, we demonstrate the utility of InducTn-seq using *Citrobacter rodentium* infection of mice, an animal model of infectious colitis. By inducing mutagenesis after a highly restrictive host bottleneck, InducTn-seq quantifies the impact of thousands of genes on bacterial fitness during enteric colonization. These analyses shed new light on the metabolic trade-offs that facilitate the transmission of diarrhea-causing bacteria.

**Fig. 1.**
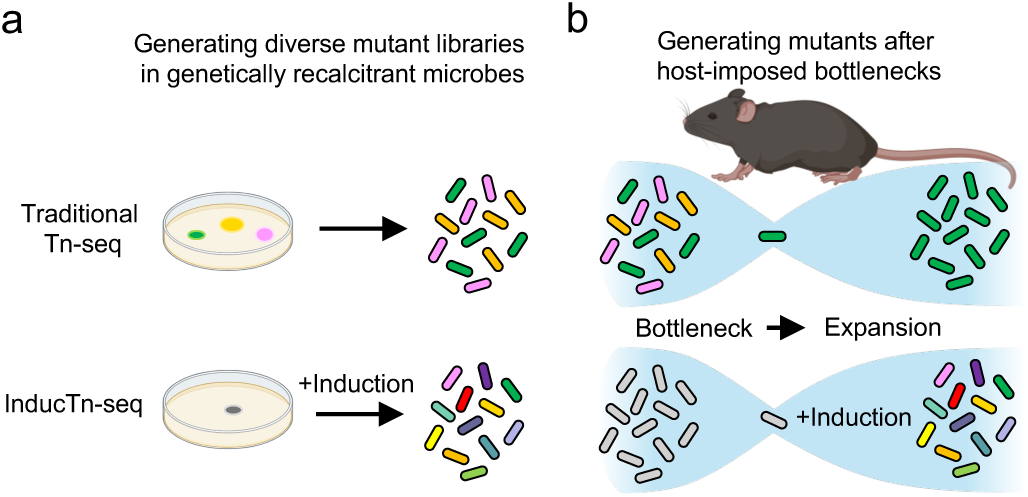
| InducTn-seq enables genome-scale forward genetics through inducible mutagenesis. **a**, Traditional Tn-seq is often limited by the inefficient delivery of a transposon donor, which impedes the generation of a diverse mutant library. By contrast, InducTn-seq allows for the expansion of a single initial transconjugant (depicted as a gray colony) to generate a diverse mutant library. **b**, During animal colonization experiments, an initial host bottleneck leads to the random elimination of mutant cells, thereby reducing the diversity of the mutant library. With InducTn-seq, new transposon mutants can be generated within the animal after bypassing the initial bottleneck.

### Design of an inducible mutagenesis system

To overcome the limitations of traditional Tn-seq, we engineered a mobilizable plasmid that introduces the randomly integrating Tn5 transposon at the *att*Tn7 site in the bacterial genome (pTn donor, Fig. 2a). The Tn5 transposon consists of a kanamycin-selectable marker flanked by hyperactive Tn5 mosaic ends^37^ and is positioned next to its corresponding Tn5 transposase gene, which is regulated by the arabinose-responsive PBAD promoter^37,38^. This Tn5 transposition complex, comprised of the Tn5 transposon and transposase sequences, is nested within the left and right ends of the *att*Tn7 site-specific Tn7 transposon^39^.

**Fig. 2.**
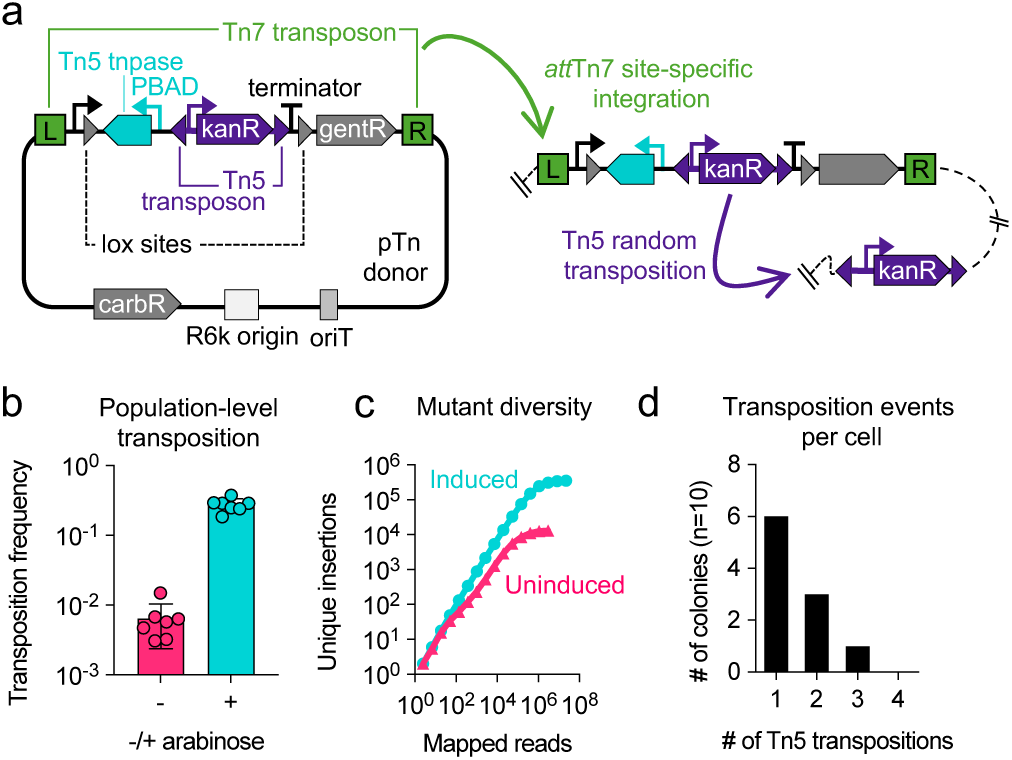
| Design of an inducible mutagenesis system. **a**, Diagram of the inducible transposon mutagenesis plasmid, pTn donor. The plasmid backbone contains a carbenicillin-selectable marker (carbR), a conditional origin of replication (R6k), and an origin of transfer (oriT). The Tn7 transposon (green brackets) contains a Tn5 transposase regulated by the arabinose-responsive PBAD promoter (cyan), a Tn5 transposon with a kanamycin-selectable marker (kanR, purple), and a transcriptionally silenced gentamicin-selectable marker (gentR, gray). The Tn5 transposase and transposon form the Tn5 transposition complex, which is flanked by Cre-recognized lox sequences. Cre excision of the complex enables the measurement of the population-level transposition frequency (see Extended Data Fig. 2 for details). Following integration of the Tn7 transposon at the *att*Tn7 site (green arrow), arabinose-mediated induction of the Tn5 transposase results in random Tn5 transposition out of the *att*Tn7 site (purple arrow). **b**, The frequency of Tn5 transposition out of the Tn7 site after growth in the presence or absence of arabinose, expressed as the ratio of kanR+gentR CFU to gentR CFU (see Extended Data Fig. 2 for details). The columns represent means, the error bars represent standard deviation, and the points represent replicates. **c**, The number of unique Tn5 insertion sites in a population of ∼10^3^ *E. coli* MG1655 colonies after growth with or without arabinose (induced or uninduced, respectively). 100 ng of template DNA was used for amplification of each sequencing library. **d**, Histogram displaying the number of Tn5 transposons inserted into the genome of ten colonies that underwent at least one Tn5 transposition event after arabinose induction. The genomic coordinates of the Tn5 transposons in each colony are provided in Supplementary Table 2.

Co-introduction of pTn donor and a Tn7 helper plasmid, which encodes the proteins necessary for Tn7 integration^40^, into a recipient strain results in the integration of the Tn5 transposition complex at the *att*Tn7 site (Fig. 2a and Extended Data Fig. 1). Subsequently, culturing *att*Tn7 site-specific integrants in the presence of arabinose triggers random Tn5 transposition (Fig. 2a). We note that Tn5 functions in a “copy-paste” manner in our experiments, creating multiple copies of the transposon within a single cell’s genome, consistent with previous literature^41,42^.

We incorporated a Cre recombinase-based indicator of the population-level transposition frequency in the design of our system. This was achieved by flanking the Tn5 transposition complex with lox sequences, which separate an upstream constitutive promoter from a downstream gentamicin selectable marker preceded by a transcriptional terminator (Fig. 2a and Extended Data Fig. 2). We introduced our mutagenesis system into the K-12 *Escherichia coli* strain MG1655 through conjugation and selected for transposon integrants using kanamycin in the presence or absence of arabinose. The Cre-based indicator revealed that Tn5 transposition was ∼43-fold higher in the presence of arabinose, demonstrating that arabinose-induced transposase expression effectively drove mutagenesis (Fig. 2b). In fact, the Cre-based readout consistently revealed that ∼28% of cells induced with arabinose were Tn5 mutants, i.e., they contained a Tn5 transposon insertion at one or more sites in the genome other than *att*Tn7 (Fig. 2b).

### High-density insertional mutagenesis with InducTn-seq

To assess the diversity of Tn5 insertions following induction, we sequenced the mutant population arising from ∼10^3^ colony-forming units (CFU) collected from plates with or without arabinose. In the absence of induction, ∼1.3×10^4^ unique insertions were detected, which we attribute to low-level leakiness of the PBAD promoter. However, following induction with arabinose, ∼3.5×10^5^ unique insertions were detected (Fig. 2c). These findings corroborate the Cre-based readout and provide additional evidence that arabinose induces mutagenesis. Interestingly, when mutagenesis is induced during colony outgrowth, the number of unique insertions in the induced population exceeds the number of colonies, indicating that each colony is comprised of a mosaic of unique mutants.

We found that the number of detectable unique insertions was constrained by the quantity of template DNA used in the preparation of the sequencing library (see Methods). Increasing the amount of template DNA sampled led to an increase in the number of unique insertions identified (Extended Data Fig. 3). With our inducible mutagenesis system, it is possible to generate a diverse transposon mutant library containing over 1 million unique insertions from a small patch of cells scraped from a single petri dish.

Inducible mutagenesis introduces the potential for multiple Tn5 insertions within a single cell. To quantify this possibility, we performed whole-genome sequencing of ten colonies, each originating from a cell that had undergone at least one Tn5 transposition event (see Methods). This analysis revealed that the majority of mutant cells in the induced population contained a single Tn5 insertion after overnight culture, with the likelihood of having additional insertions within the same cell decreasing exponentially, consistent with a Poisson distribution (Fig. 2d, Supplementary Table 2, and Discussion).

### Insertions in essential genes

In contrast to traditional Tn-seq, gene-level analysis of transposon insertions revealed that virtually all genes were heavily mutagenized in the induced population, with minimal difference in insertion frequency between canonically essential and non-essential genes. For example, the genes *obgE*, *rpmA*, *rplU*, *ispB*, and *murA*, which are canonically essential in *E. coli*^43,44^, exhibited a similar number of insertions in the induced population as the neighboring non-essential genes (Fig. 3a). This suggests that the rate of mutagenesis during induction surpasses the rate of selection against transposon insertions in essential genes.

**Fig. 3.**
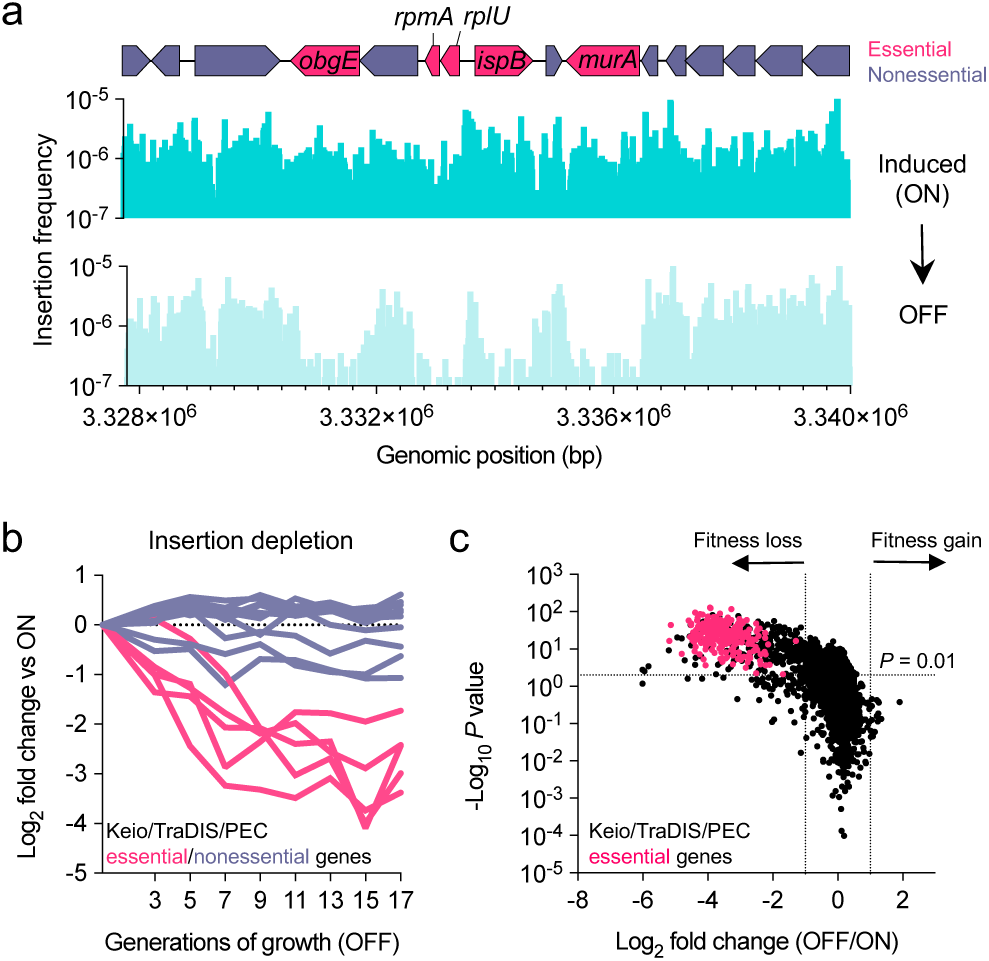
| Sensitive measurement of mutant fitness in both essential and non-essential genes. **a**, Induced cells (ON) contain Tn5 insertions in genes classified as essential in the closely related *E. coli* strain BW25113 (genes depicted in red, e.g., *obgE*). Insertions in essential genes are selectively depleted when the population is expanded in the absence of induction (**a**), and progressively decrease with more generations of growth (**b**). In **b**, the ON population was serially diluted in LB without induction, ensuring logarithmic growth of the population over 17 generations. Individual lines in panel **b** correspond to the genes displayed in panel **a**. See Supplementary Table 3 for a complete list of all genes. **c**, Volcano plot comparing the fold change in the insertion frequency between OFF and ON. A significant fitness defect was defined as a Mann Whitney U *P* value <0.01 and log_2_ fold change <-1 relative to the frequency of insertions in the ON population. Genes previously classified as essential are marked as red points.

We hypothesized that further culturing the induced population in the absence of arabinose would lead to the selective depletion of transposon insertions in essential genes. To test this hypothesis, we continually passaged the induced population under non-inducing conditions and found that the magnitude of depletion increased with successive generations (Fig. 3 a-b and Supplementary Table 3). Based on these findings, we refer to the induced population as “ON” and further outgrowth in the absence of induction as “OFF”. The transition from ON to OFF allows us to observe the effects of selection on the mutant population.

### A sensitive, streamlined framework for mutant fitness analysis

A direct comparison of the insertion frequency within each gene between the ON and OFF populations could improve the analysis of mutant fitness for both essential and non-essential genes. In traditional Tn-seq, genes are classified as either essential or non-essential based on their insertion frequency relative to the genome-wide insertion density. However, this approach is susceptible to both false positive and false negative essential gene calls, especially in shorter, AT-rich genes and in libraries with low diversity^5^. By directly comparing each gene to itself between the ON and OFF conditions, InducTn-seq controls for these confounding factors in data analysis. This simplified approach to essential gene analysis is fundamentally unattainable with traditional Tn-seq outside of organisms with exceptionally high transformation and recombination efficiencies^45^, because insertions in essential genes are absent within the initial library.

We assessed the accuracy of InducTn-seq using this analytic framework by comparing the insertion frequency in each gene between the ON and OFF conditions. We classified a gene as having a fitness defect if it exhibited more than a two-fold reduction in its insertion frequency (log_2_ fold change <-1) and a corrected *P*-value <0.01 when comparing OFF to ON (see Methods). With these criteria, 532 genes out of 4,494 annotated gene features in the *E. coli* MG1655 genome (NCBI Reference Sequence: NC_000913.3) had a fitness defect after dense overnight growth on solid LB (Fig. 3c and Supplementary Table 4).

We compared our results to a set of 248 genes consistently classified as “essential” across three datasets in the closely related *E. coli* strain BW25113^44,46,47^. Our screen identified a fitness defect in all 248 genes, demonstrating its sensitivity (Fig. 3c and Supplementary Table 5). Among the three comparator datasets, two originated from targeted gene knockout studies (Keio and PEC), while the third dataset was derived from a traditional Tn-seq experiment (TraDIS)^44,46,47^. To gauge the performance of InducTn-seq relative to traditional Tn-seq, we directly compared our dataset to the TraDIS dataset, which represents one of the most highly saturated transposon mutant libraries generated in *E. coli*, prior to this study^44^. In this comparison, InducTn-seq identified 331 of the 354 genes classified as essential by TraDIS and annotated in the MG1655 genome, a 93.5% overlap (Extended Data Fig. 4a and Supplementary Table 6). Importantly, the 23 genes identified solely by TraDIS were also not identified as essential in either the Keio or PEC datasets, which are predominantly based on gene information gathered via genomic deletion or allelic exchange rather than transposon insertion. To reconcile the discrepancies between our dataset and the TraDIS dataset, we quantified the gene length, GC content, and fold change of genes with fitness defects identified by the two screens. We discovered that the 23 genes identified by TraDIS, but not by InducTn-seq, Keio, or PEC, were significantly shorter and more AT-rich, with a median gene length of 204 bp and median GC content of 32.7%, compared to a median of 879 bp and 52.8% among genes identified by both Tn-seq based methods (Extended Data Fig. 4b,c and Supplementary Table 7). These data suggest that many of the genes identified solely by TraDIS may be false positives due to the lower likelihood of transposon insertions in shorter, AT-rich genes^48–51^.

Furthermore, we found that the 201 genes identified by InducTn-seq, but not by TraDIS, had a median log_2_ fold change of –1.86 compared to a median of –3.44 among genes identified by both methods (Extended Data Fig. 4d and Supplementary Table 7). This implies that the genes identified solely by InducTn-seq may have relatively weaker fitness defects that fall below the cutoff of being classified as “essential” by TraDIS analysis.

Collectively, our results demonstrate that InducTn-seq accurately identifies known essential genes, is exceptionally sensitive for detecting genes with more subtle fitness defects, and is robust to false positives in short, AT-rich genes. Moreover, by having an ON to OFF comparator, InducTn-seq transforms the traditionally binary classification of “essential” vs. “nonessential” into a quantitative measurement of fitness^52^.

### High-diversity mutant libraries originating from a single colony

We next assessed the versatility of InducTn-seq outside of K-12 *E. coli* by generating mutant libraries in four enteric pathogens (*Citrobacter rodentium*, Enterotoxigenic *E. coli*, *Salmonella enterica* serovar Typhimurium, and *Shigella flexneri*). We introduced pTn donor through conjugation and selected for transposon integrants. Although these pathogens are all related gammaproteobacteria, there was substantial variation in transconjugant frequency, spanning nearly four orders of magnitude across species (Fig. 4a).

**Fig. 4.**
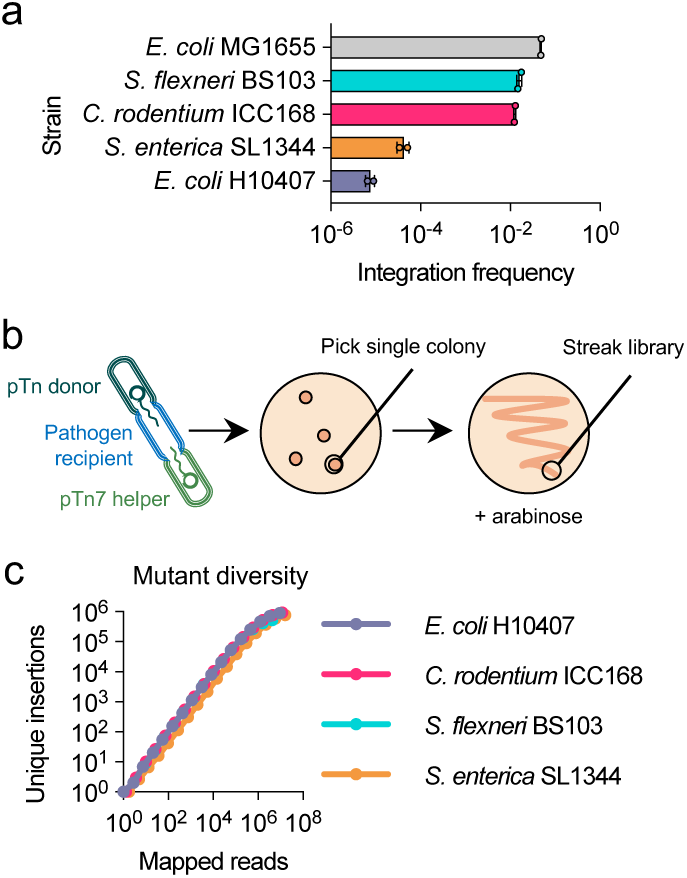
| InducTn-seq creates high-density mutant libraries from a single bacterium. **a**, The indicated enteric pathogens underwent conjugation with pTn donor and pTn7 helper. The integration frequency is expressed as the ratio of kanR CFU to total CFU. The bars represent means and the error bars represent standard deviation. n = 2 for each strain. **b**, A single, uninduced transconjugant colony of each strain was streaked onto a plate containing 0.2% arabinose. **c**, The number of unique Tn5 insertion sites detected by sequencing of the streaked libraries expanded from the single colony.

Creating a diverse mutant population using traditional mutagenesis is often hindered by low efficiency of transposon DNA delivery to recipient cells. However, InducTn-seq theoretically requires only a single initial integration event at the *att*Tn7 site to enable subsequent generation of a mutant population with induction. To test the power of inducible mutagenesis and mimic the bottleneck these pathogens might encounter during animal colonization, we experimentally bottlenecked the population by picking a single transconjugant colony of each strain and streaking it onto a new plate with arabinose to generate the ON population (Fig. 4b). Starting with just a single colony, we were able to create transposon mutant populations in each pathogen with 10^5^-10^6^ unique insertions (Fig. 4c). Taken together, these findings underscore the broad applicability of InducTn-seq across species and its capacity to overcome bottlenecks that reduce the population to a single cell.

### Using InducTn-seq to circumvent the host bottleneck

*C. rodentium* is commonly used as a murine model of infectious colitis because it shares an infection strategy with the attaching and effacing (A/E) family of human extracellular pathogens. All A/E pathogens encode the conserved Locus of Enterocyte Effacement (LEE) pathogenicity island^53,54^, but in contrast to human A/E pathogens (EHEC and EPEC), *C. rodentium* can infect and cause diarrhea in mice without the need for antibiotic sensitization. However, a severe bottleneck impedes *C. rodentium* intestinal colonization, where only about 10^1^-10^2^ unique cells initiate a typical infection^31,32^. Given that genome-scale Tn-seq libraries usually contain >10^5^ unique mutants, traditional Tn-seq is not feasible in this model due to the random elimination of most mutants by the restrictive bottleneck.

We hypothesized that inducing mutagenesis after the bacterial population underwent a bottleneck would enable identification of the genetic requirements for *C. rodentium* colonization. To test this hypothesis, we performed InducTn-seq in C57BL/6J mice and compared it to traditional Tn-seq. For the traditional method, mice were inoculated with ∼10^10^ cells comprising ∼3×10^5^ unique *C. rodentium* transposon mutants. For InducTn-seq, we administered a similar number of uninduced cells and induced mutagenesis after the severe constriction of the *C. rodentium* mutant population by the host/microbiota bottleneck^32^. Mutagenesis was induced by providing mice with 5% arabinose in their drinking water from days 3 to 8. The *C. rodentium* population was tracked by enumerating fecal CFU, and notably, arabinose induction did not affect the abundance of fecal *C. rodentium* (Fig. 5a).

**Fig. 5.**
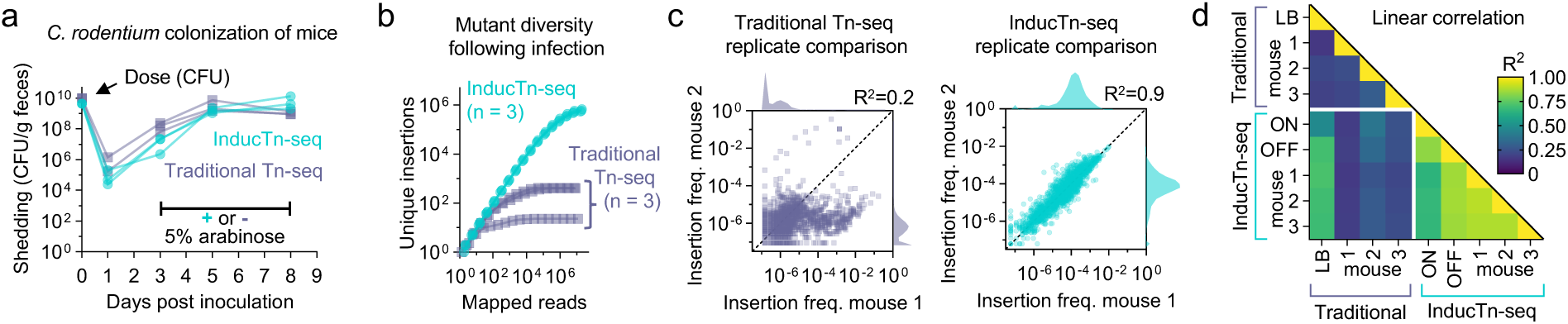
| InducTn-seq bypasses the host bottleneck. **a**, Female C57BL/6J mice were intragastrically inoculated with either a pool of ∼3×10^5^ unique *C. rodentium* Tn5 insertion mutants (Traditional Tn-seq) or uninduced Tn5 transposition complex integrants (InducTn-seq), and colonization was monitored by serial dilution and plating of feces. For InducTn-seq, Tn5 transposition was induced on day 3 to 8 by providing *ad libitum* access to water containing 5% arabinose. n = 3 mice in each group. **b**, Samples from day 5 (Traditional Tn-seq) or day 8 (InducTn-seq) post-inoculation were sequenced to determine the number of unique mutants recovered from each animal. n = 3 mice in each group. **c**, Correlation of mutant frequency between animal replicates. Points represent genes, insertion frequency is calculated as reads per gene normalized to total reads in the sample, and histograms on the axes display the distribution of the data. **d**, The coefficient of determination (R^2^) comparing the log_10_ transformed insertion frequencies across replicates.

As expected, the diversity of the traditional Tn-seq library was dramatically reduced during infection, resulting in a population of ∼10^2^ unique mutants 5 days post-inoculation (Fig. 5b). Because of this random loss of mutants, the identity and abundance of recovered mutants varied markedly between animals (average R^2^ = 0.24, Fig. 5c,d). In stark contrast, induction of mutagenesis *in vivo* resulted in a highly diverse mutant library within the infected animals, allowing recovery of >5×10^5^ unique mutants 8 days post-inoculation (Fig. 5b), with consistent gene-level insertion frequencies between animals (average R^2^ = 0.89, Fig. 5c,d). Thus, InducTn-seq resolved the bottleneck problem that prevents traditional *in vivo* transposon mutant screens.

### The benefits of plasticity in oxygen-related metabolism

Using our new analytic framework, we identified the genes that contribute to *C. rodentium* fitness during animal colonization by comparing the mutants recovered from the feces of an infected mouse (mouse-OFF) to a population induced on solid LB containing arabinose (LB-ON) (schematized in Fig. 6a and displayed on the x-axis of 6b, see also Supplementary Table 8). We also determined the genes required for *C. rodentium* fitness in culture by comparing cells expanded on solid LB in the absence of arabinose induction (LB-OFF) to LB-ON (displayed on the y-axis of 6b).

**Fig. 6.**
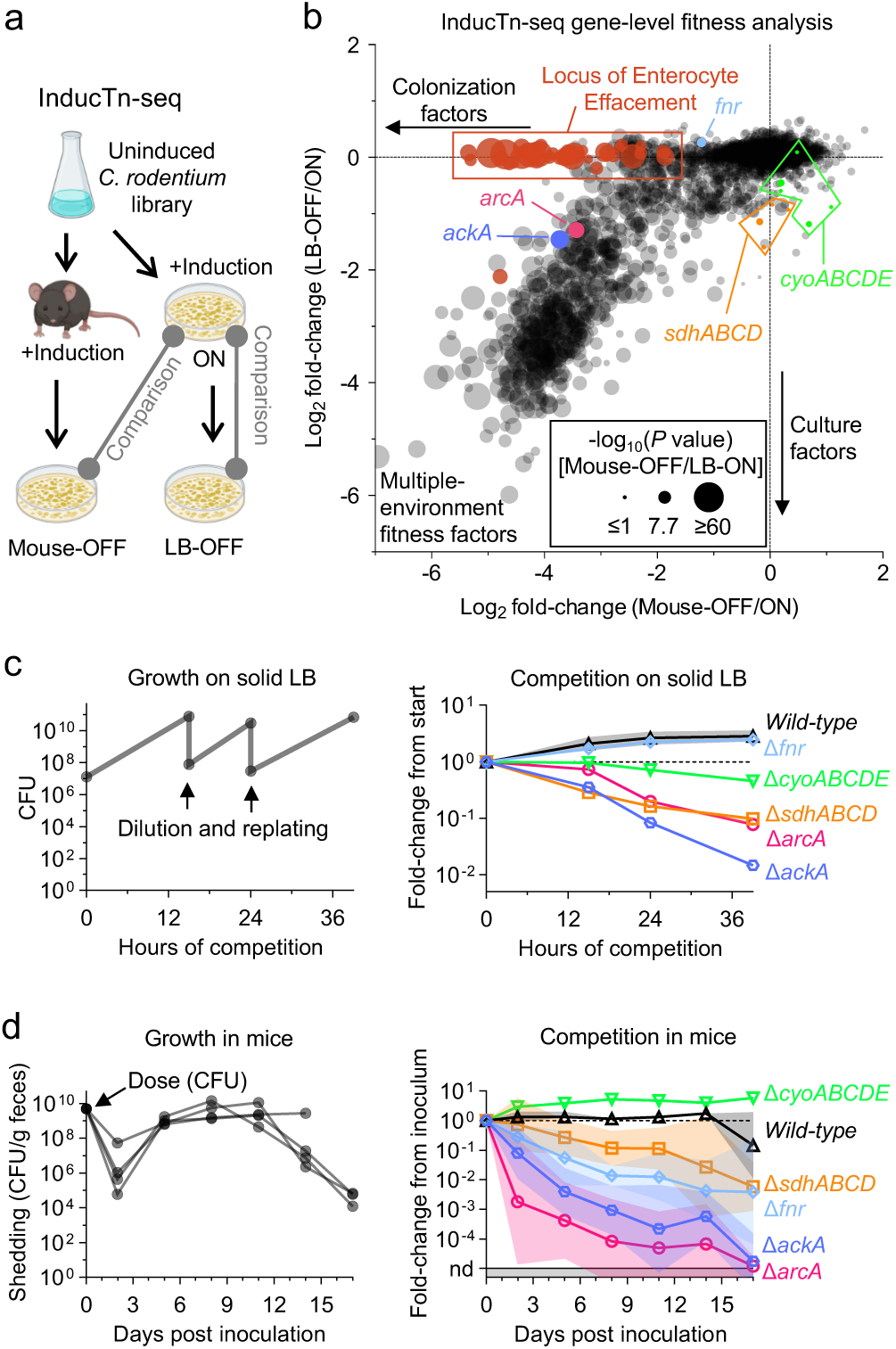
| InducTn-seq reveals the role of oxygen-related metabolism during enteric infection. **a**, Experimental scheme. Mutant populations isolated from the mouse (mouse-OFF) or following *in vitro* outgrowth in the absence of induction (LB-OFF) were each compared to the induced population generated *in vitro* (LB-ON). **b**, Fold change in the insertion frequency during infection (mouse-OFF) or during culture (LB-OFF) relative to induction (LB-ON), as schematized in **a**. Points represent genes and significance represents a Mann-Whitney U test comparing mouse-OFF to LB-ON. **c-d,** To validate the InducTn-seq results, barcoded strains with the indicated in-frame deletions were expanded separately and then pooled and either passaged on LB agar plates (**c**) or intragastrically administered to female C57BL/6J mice (**d**). The size of the bacterial population was determined by serial dilution and plating, while the relative abundance of each strain was measured by amplicon sequencing. Lines represent geometric means and error bars represent standard deviations of three barcoded strains per deletion competed in a single culture (**c**) or four barcoded strains per deletion competed in four mice (**d**). ND = barcode not detected.

Mutants with a fitness defect in mice included previously identified *C. rodentium* colonization factors. For example, multiple studies have shown that the LEE pathogenicity island is required for *C. rodentium* to colonize mice^53–57^. In line with these findings, fewer insertion mutants in LEE genes were recovered from mouse-OFF than LB-ON (x-axis, Fig. 6b). In contrast, expanding the mutants generated in culture without induction (LB-OFF) did not decrease the frequency of mutants in LEE genes (y-axis, Fig. 6b), indicating that the LEE’s function is required during infection, but not during culture (i.e., a colonization-specific factor). InducTn-seq also identified other previously characterized *C. rodentium* colonization factors, including the virulence gene regulator RegA^58^, multiple amino acid biosynthesis pathways^23^, and enzymes involved in aerobic respiration^59^ (Supplementary Table 8). These results demonstrate that InducTn-seq enables reproducible genome-scale forward genetics in an animal model of infection with a highly restrictive colonization bottleneck.

Like most diarrheal pathogens (e.g., Shigella, Vibrio, Salmonella, Listeria, and Escherichia spp.), *C. rodentium* thrives in both the oxygen-poor environment of the intestine and the oxygen-rich environment outside of the host (i.e., it is a facultative anaerobe). We observed that the genes enabling this metabolic plasticity in oxygen utilization exhibited a range in phenotypes, from culture-specific (e.g. *cyoABCDE*) to animal-specific (e.g. *fnr*) (highlighted in Fig. 6b).

We focused on five key genes/complexes: (1) *ackA*, active in both anaerobic and aerobic acetate production; (2) *fnr*, a transcriptional regulator active exclusively in anaerobic conditions; (3) *arcAB*, a two-component system active in microaerobic conditions; and (4-5) the operons *sdhABCD* and *cyoABCDE*, which are active in the TCA cycle and the electron transport chain (ETC) in fully aerobic conditions^60–65^. InducTn-seq revealed that Fnr-dependent anaerobic metabolism provides *C. rodentium* with a specific advantage during animal colonization (a colonization-specific factor), while ArcAB-dependent microaerobic metabolism is important for fitness during both animal colonization and growth in culture (a multiple environment fitness factor). In contrast, fully aerobic metabolism confers a fitness advantage exclusively outside of the host (a culture-specific factor) but impairs the pathogen’s fitness within the host (*i.e.,* insertion mutants in *cyoABCDE* become more prevalent in infected mice).

To validate the InducTn-seq data, we engineered *C. rodentium* strains with in-frame deletions in *ackA*, *fnr*, *arcA*, and the *sdh* and *cyo* operons (*sdhABCD* and *cyoABCDE*). We introduced unique nucleotide sequences as barcodes into each strain at the *att*Tn7 site^27^. These ∼25 bp barcodes are identifiable by amplicon sequencing and enable us to monitor the relative abundance of each strain in pooled competition experiments.

These mutant strains were cultured separately, and then pooled with wild-type *C. rodentium* and serially passaged on solid LB (Fig. 6c, left). Consistent with our InducTn-seq results (Fig. 6b), we observed that strains lacking genes involved in general (Δ*ackA*), microaerobic (Δ*arcA*), and fully aerobic (Δ*sdhABCD* and Δ*cyoABCDE*) metabolism were outcompeted within the pool. In contrast, strains lacking the master transcriptional regulator of anaerobic metabolism (Δ*fnr*) did not exhibit a fitness defect in aerobic culture (Fig. 6c, right).

This pool of strains was also intragastrically inoculated into mice and their relative abundance was monitored by plating feces from colonized animals (Fig. 6d, left). As indicated by InducTn-seq (Fig. 6b), deletion of genes involved in general (Δ*ackA*), anaerobic (Δ*fnr*), and microaerobic (Δ*arcA*) metabolism impaired the pathogen’s fitness during intestinal colonization (Fig. 6d, right). Deletion of *sdhABCD* also caused a minor colonization defect not detected by InducTn-seq, a discrepancy that could be attributable to the differences in the time points of infection assayed in the barcoded competition (i.e., *sdh* genes may have a role early during infection before induction, which began on day 3). Intriguingly, strains lacking *cyoABCDE* had a ∼6-fold growth advantage inside of the animal (Fig. 6d, right), consistent with our InducTn-seq results and suggesting that this complex is detrimental to the pathogen’s fitness during animal colonization.

CyoABCDE, a low-affinity terminal oxidase, is used in the ETC exclusively in oxygen-rich environments^65–68^. Notably, obligate anaerobic commensal bacteria, which constitute the majority of gut microbes^69–71^, do not encode this complex. The results revealed by InducTn-seq could partially account for the contrasting metabolic strategies observed in pathogenic and non-pathogenic bacteria. Pathogens, which frequently transmit between hosts, spend a larger proportion of their lifecycle outside of the intestine compared to commensals, which generally cannot survive for long in atmospheric oxygen. We propose that genes such as *cyoABCDE*, which are specialized for promoting growth in oxygenated non-host environments, provide a net benefit to pathogens by facilitating transmission, despite their fitness cost during infection. These findings underscore the metabolic tradeoffs inherent to the enteric pathogen life cycle, which must balance growth inside and outside of the host.

## Discussion

Despite its inherent limitations, Tn-seq is the method of choice for genome-scale forward genetics in bacteria because of its simplicity, cost-effectiveness, and power for uncovering the relationship between genotype and phenotype. InducTn-seq retains these advantages while creating new avenues for discovery by enabling the generation of mutants on demand. The ability to control mutagenesis temporally and with high efficiency offers several benefits over traditional, one-time transposition, including: (1) the ability to generate millions of mutants from a single clone, thereby overcoming barriers to conjugation and transformation in genetically recalcitrant microbes, (2) the creation of unparalleled diversity, which enhances the sensitivity of detecting subtle phenotypes, (3) the capacity to quantify the fitness impact of canonically essential genes, moving beyond a binary classification of essentiality, and (4) the ability to induce mutagenesis after host-imposed colonization bottlenecks, permitting functional genetics in new animal models of infection. These advantages will empower functional genetics in novel species in both ecologically and biomedically relevant conditions.

Historically, inducible mutagenesis has been avoided in functional genetic screens, presumably to prevent the accumulation of multiple insertions within the same cell. Multiple transposon insertions within a clone can obscure the causal relationship between an insertion and an observed phenotype and introduces the possibility of a phenotype arising due to a genetic interaction between insertions. The presence of a single, stably integrated transposon is crucial when isolating clones for phenotypic characterization or when creating an ordered transposon mutant library. However, pooled screening of transposon mutants using Tn-seq is robust against the effects of rare genetic interactions in cells with multiple transposon insertions because the fitness of a gene knockout is determined by the collective behavior of tens to hundreds of unique insertions in a high-density mutant population. Additionally, the statistical tests used to analyze Tn-seq data^5^ rely on information from multiple unique sites, buffering against the effects of rare genetic interactions within the same cell. Consequently, inducible mutagenesis can be used in Tn-seq experiments with minimal risk of multiple insertions in the same cell confounding the results.

While our attempt to perform a genome-scale screen in mice using a traditional *C. rodentium* mutant library was unsuccessful (Fig. 5), Caballero-Flores *et. al*. previously used a traditional Tn-seq approach to identify genes required for *C. rodentium* colonization^23^. Since the bottleneck to *C*. *rodentium* colonization is primarily dependent on the microbiota^32^, the more permissive bottleneck observed in the Caballero-Flores *et. al.* screen is likely due to variations in the composition of the murine microbiota between institutions. Consistent with this idea, our analyses of the gene-level insertion data from Caballero-Flores *et. al.* revealed much more variability between mice with a conventional microbiota (specific pathogen free, SPF) than germ-free animals, which lack a microbiota (SPF R^2^ = 0.50 vs. germ-free R^2^ = 0.99, Extended Data Fig. 5). Various treatments, including antacids to neutralize gastric acid or broad-spectrum antibiotics to deplete the commensal microbiota^23,31,32,72^, are often used to widen the host bottleneck when implementing traditional Tn-seq in mouse models of intestinal colonization. While using germ-free mice or one of these treatments may bypass the bottleneck problem, these interventions also disrupt the natural dynamics between the host and microbe, altering the bacterial genetic requirements for colonization. For example, the LEE is required for *C. rodentium* to colonize SPF mice (Fig. 6b and Supplementary Table 8), but is dispensable in germ-free animals^73^. One of the major advantages of InducTn-seq over traditional approaches is that it facilitates performing bacterial genetic screens in experimental systems without the need to characterize or disrupt natural colonization bottlenecks.

Another advantage of InducTn-seq in animal studies is its sensitivity. Since we recovered >10^5^ unique *C. rodentium* mutants from infected mice, we were able to resolve subtle phenotypes and uncover metabolic pathways that facilitate pathogenesis. We found that Fnr-dependent anaerobic metabolism benefited *C. rodentium* within the animal without causing a fitness defect in oxygenated environments outside of the host. In contrast, the genes (*cyoABCDE*) used for metabolism in highly oxygenated environments provide a subtle fitness advantage in culture at the cost of a subtle fitness disadvantage during infection (Fig. 6b-d). The ability of InducTn-seq to quantify these metabolic trade-offs provides valuable insight into the life cycle of enteric pathogens and has the potential to open novel avenues for understanding the evolution of bacterial pathogenesis. Moreover, leveraging inducible mutagenesis in animal-based studies will facilitate the spatiotemporal characterization of genetic programs employed by pathogens during different stages of infection and in different tissues.

Although we used the arabinose-responsive PBAD promoter to induce transposase expression^38^, the concept of inducible transposition can be extended to other regulatable promoter systems. For example, control of mutagenesis could be achieved using the anhydrotetracycline-responsive TetR system, which is appealing due to its bio-orthogonality^74^. Further, a temperature-inducible promoter active at 37 °C could be particularly valuable for inducing transposase expression specifically in mammalian hosts because it would eliminate the need to supply an inducer molecule^75,76^. While Tn7-based integration will not be applicable in many microorganisms, additional site-specific integration systems, such as phage-based integration vectors for *Mycobacterium tuberculosis* and *Listeria monocytogenes*^77,78^, should be useful for chromosomal integration of an inducible transposition system. We also note that InducTn-seq is effective when expressed from a replicative plasmid rather than being integrated into the genome, making it applicable in organisms that lack the *att*Tn7 site or other means of genomic integration. In summary, InducTn-seq should be widely adaptable to diverse microbial systems, including non-prokaryotic systems such as the human parasite *Plasmodium falciparum*, where Tn-seq has previously been applied^79^.

## Methods

### Bacterial strains

The strains used in this manuscript are listed in Supplementary Table 1. For cloning, we used the *E. coli* strain MFD*pir*^80^. Mutant libraries were made in the K-12 *E. coli* strain MG1655, Enterotoxigenic *E. coli* (ETEC) strain H10407, *Salmonella enterica* serovar Typhimurium strain SL1344, and *Shigella flexneri* strain BS103 (a derivative of strain 2457T lacking the virulence plasmid). For animal infection experiments we used a spontaneous streptomycin-resistant isolate of *Citrobacter rodentium* strain ICC168^32^.

### Plasmid assembly

Assembly of the transposon mutagenesis plasmid pTn donor (pTn) and the Cre-expressing plasmid pCre was performed using the NEBuilder HiFi DNA Assembly Master Mix (New England Biolabs). Fragments were generated through PCR amplification from plasmid or genomic DNA templates using primers acquired from Integrated DNA Technologies (IDT). The ribosome binding site (RBS) located upstream of the Tn5 transposase open reading frame (ORF) in pTn was designed using the RBS calculator v2.2^81–83^. The target translation initiation rate was set at 20,000 arbitrary units. The *cre* ORF in pCre was amplified from an arabinose-regulated Cre-expression vector^84^. Following assembly, the plasmids were cloned into the *E. coli* strain MFD*pir* using electroporation. Each plasmid sequence was then verified through whole-plasmid sequencing (Primordium Labs). All plasmid sequences were edited using the open source software “A plasmid Editor” (ApE)^85^. The plasmid sequences were annotated using pLannotate^86^ and are provided as supplementary files.

### Generation of induced transposon mutant populations

Induced mutant populations were generated using the donor strain MFD*pir*+pTn and the helper strain MFD*pir*+pJMP1039, which expresses the Tn7 transposase enzymes^40^. Both strains were cultured in liquid LB containing 1 mM diaminopimelic acid (DAP) and 50 µg/ml carbenicillin, while recipient strains were cultured in liquid LB without DAP or antibiotics. The cultures were incubated overnight at 37 °C with shaking (250 rpm).

The following day, 1 ml of each overnight culture was washed once with 1 ml of fresh LB, and then resuspended in 50-100 µl of LB. The cultures were mixed in a 1:1:1 ratio of MFD*pir*+pTn, MFD*pir*+pJMP1039 helper, and recipient strain. A 60 µl aliquot of this mating mixture was spotted onto a 0.45 µm pore mixed cellulose ester filter disc (Millipore) placed on solid LB containing DAP and incubated at 37 °C for 2 h.

After incubation, the filter discs were removed from the plate using sterile forceps and resuspended in 1 ml of LB in a microfuge tube by vortexing. The resuspended mating mixtures were then diluted and plated on solid LB containing 50 µg/ml kanamycin and 0.2% L-arabinose to induce Tn5 transposase expression. For the *E. coli* H10407 recipient strain, 100 µg/ml of kanamycin was used. After overnight growth, colonies were scraped from the plates, resuspended in liquid LB containing 15% glycerol, and stored as aliquots at –80 °C.

To induce mutant populations of enteric pathogens starting from a single colony, resuspended mating mixtures were first plated on solid LB containing kanamycin without arabinose. After overnight growth, a single transconjugant colony of each strain was then streaked onto solid LB containing kanamycin and arabinose to form the induced mutant population.

### Assessment of Tn5 and Tn7 integration frequency

The integration frequencies of Tn5 and Tn7 were assessed by serial dilution and plating of the resuspended mating mixtures on solid LB with our without kanamycin to enumerate colony-forming units (CFU)^87^. The integration frequency, representing the combined total of Tn5 and Tn7 integration, was quantified by comparing the number of kanamycin-resistant CFU to total CFU. To assess the frequency of Tn5 random integration alone, MFD*pir*+pJMP1039 (Tn7 helper plasmid) was excluded from the mating mixture.

### Assessment of Tn5 transposition frequency and transposition events per cell

The transposition frequency of Tn5 was assessed by plating the resuspended mating mixtures on solid LB containing 50 µg/ml kanamycin, with or without 0.2% L-arabinose. After overnight growth, colonies were scraped from each plate and a second conjugation was performed with MFD*pir*+pCre. Following this second conjugation, the resuspended mixtures were serially diluted and plated on solid LB containing 20 µg/ml gentamicin with or without 50 µg/ml kanamycin. The transposition frequency was quantified by comparing the number of gentamicin and kanamycin-resistant CFU to gentamicin-resistant CFU (Extended Data Fig. 2).

Cre activity excises the Tn5 transposase, thereby preventing further mutagenesis and leaving the cell with a fixed number of Tn5 insertions. To determine the number of transposition events per cell following Cre recombination, ten gentamicin and kanamycin-resistant colonies (i.e., originating from cells that had undergone at least one Tn5 transposition event) were individually streaked onto solid LB and sent to the Microbial Genome Sequencing Center (Pittsburgh, PA) for whole genome sequencing. Among ten sequenced colonies, we found that six had a single Tn5 insertion, three had two insertions, and one had three insertions (Fig. 2d and Supplementary Table 2).

### Assessment of insertion depletion in the absence of induction

The depletion of transposon insertions was assessed by performing repeated serial dilution and outgrowth of the ON population in the absence of induction (OFF). A 50 µl aliquot of a frozen glycerol stock of the *E. coli* MG1655 ON population was diluted into 5 ml of LB to achieve a starting concentration of 5×10^7^ CFU/ml and incubated at 37 °C with shaking at 250 rpm.

After 2 h of incubation, a 2 ml aliquot of the culture was pelleted for sequencing library preparation and serial dilutions were performed to enumerate CFU. The culture was then back-diluted four-fold into fresh LB and incubated for another hour. This growth and dilution step was repeated hourly for a total of 17 generations of growth, which is approximately two generations of growth per hour. An aliquot was pelleted and CFU were enumerated at each time point.

### Sequencing library preparation

Sequencing libraries were prepared by modifying established protocols reviewed in Opijnen & Camilli^88^. In brief, genomic DNA was extracted from pelleted aliquots of the mutant populations using the Qiagen DNeasy Blood and Tissue Kit. The extracted DNA was then sheared to a size of 400 bp using a Covaris M220 ultrasonicator. The sheared DNA was end repaired using the NEB Quick Blunting kit, followed by the addition of Poly(A) nucleotide overhangs using Taq polymerase and dATP. Hybridized Illumina P7 adapters with complementary “T” overhangs were then ligated to the A-tailed DNA using NEB T4 ligase.

The end of the Tn5 transposon is flanked by cut sites recognized by the restriction enzymes PacI and SpeI (see annotated plasmid sequence in Supplementary File 1). After adapter ligation, the DNA was digested with PacI and SpeI, resulting in the creation of an ∼90 bp fragment containing the transposon end within the integrated vector sequence. Following digestion, the DNA was size-selected using SPRIselect beads at a 1× ratio to remove the digested fragment. This step obviates the need to cure the integrated portion of the plasmid vector, streamlining the generation of both transposon mutant and sequencing libraries, and eliminating uninformative sequencing reads that map to the integrated vector.

Following size selection, 100-800 ng of the DNA was amplified for 20 cycles using a transposon-specific forward primer and an adapter-specific reverse primer. Each reverse primer contained a unique Illumina i7 index sequence. The PCR products underwent a second round of 1x size selection to remove any primer dimer. The concentration of the sequencing libraries was quantified using a Qubit fluorimeter and then pooled and sequenced on a Nextseq 1000. The sequencing reads are deposited in the sequencing read archive (SRA) under the accession number PRJNA1113708.

### Tn-seq data analysis

Sequencing reads were trimmed of the transposon end sequence and mapped to the reference genome using the Burrows-Wheeler Aligner (BWA) software package^89^ in Python. Mapped reads were then converted into a table containing a tally of the read count at each genomic position and the corresponding annotated gene name at that position. The counts were normalized for read depth and the log_2_ fold change in insertion frequency was calculated for each gene feature by comparing OFF to ON. To determine statistical significance, the genome-wide counts for each sample were first summed within a window size set as the inverse of the frequency of non-zero positions for the sample with fewer overall non-zero positions. Setting this window size helps control for differences in the overall diversity of transposon insertions between two samples. Following summation of the counts into equal window sizes, the non-parametric Mann-Whitney U statistical test was performed, and *P* values were adjusted for multiple testing using the Benjamini-Hochberg correction.

To visualize the number of unique insertion sites per sample across different sampling depths, the table of read counts per genomic position was filtered by removing the bottom 1% of all reads. This step was performed to remove noise resulting from Illumina index sequence hopping. After this noise correction, the distribution of counts per site was subjected to multinomial resampling beginning with the maximum read count of each sample and subsampling in two-fold lower increments. At each sampling depth, the number of unique insertions was calculated.

The following genomic reference sequences were used for read mapping: *E. coli* MG1655 (NC_000913.3), *E. coli* H10407 (NC_017633.1), *S. flexneri* 2457T (NC_004741.1), *S. enterica* SL1344 (FQ312003.1), and *C. rodentium* ICC168 (FN543502.1).

### Mouse experiments

Animal studies were conducted at the Brigham and Women’s Hospital in compliance with the Guide for the Care and Use of Laboratory Animals and according to protocols reviewed and approved by the Brigham and Women’s Hospital Institutional Animal Care and Use Committee (protocol 2016N000416). Adult (9-12 weeks), female, C57BL/6J mice were purchased from Jackson Laboratory (strain #000664) and acclimated for at least 72 h prior to experimentation. Mice were housed in a biosafety level 2 (BSL2) facility under specific pathogen free conditions at 68-75°F, with 50% humidity, and a 12 h light/dark cycle.

### C. rodentium infections

Mice were infected by intragastric gavage with the indicated strains or mutant libraries. Prior to inoculation, mice were deprived of food for 3-5 h. Mice were then mildly sedated by inhalation of isoflurane and 100 µl of bacteria were delivered into the stomach with a sterile feeding needle (Cadence Science). The dose was determined retrospectively by serial dilution and plating.

The *C. rodentium* population was monitored by collecting feces from infected animals. Feces were weighed, resuspended in sterile phosphate-buffered saline (PBS), and homogenized using 3.2 mm stainless-steel beads and a bead beater (BioSpec Products, Inc). The concentration of *C. rodentium* was determined by serial dilution. Tn-seq and barcode libraries were expanded from feces plated on solid LB containing streptomycin and/or kanamycin and the bacteria were frozen for sample processing.

### Tn-seq of *C. rodentium* during infection

Inducible and traditional libraries were prepared by collecting ∼5×10^5^ kanamycin resistant colonies from the conjugation of *C. rodentium* to MFD*pir*+pTn with (inducible) or without (traditional) the Tn7 helper strain MFD*pir*+pJMP1039. The inducible library was cultured with 0.2% glucose prior to and following infection to prevent mutants arising outside of the animal. Aliquots of the libraries were frozen at –80 °C in 20% glycerol. Immediately prior to infection, the libraries were thawed and expanded for 3 h in LB at 37 °C with shaking at 250 rpm. The bacteria were then pelleted, resuspended in PBS, and administered to the animals by intragastric gavage.

For InducTn-seq, the animal’s drinking water was replaced from days 3-8 with water containing 5% L-arabinose (Sigma Aldrich). InducTn-seq samples were collected 8 days post inoculation by plating fresh feces on solid LB containing 50 µg/ml kanamycin, 200 µg/ml streptomycin, and 0.2% glucose. For traditional Tn-seq, the animal’s drinking water was not supplemented with arabinose and Tn-seq samples were collected 5 days post inoculation by plating feces on solid LB containing 50 µg/ml kanamycin and 200 µg/ml streptomycin.

### Constructing mutant strains of *C. rodentium*

In-frame deletions of *ackA, fnr, arcA, sdhABCD, and cyoABCDE* from *C. rodentium* were constructed using the allelic exchange protocol from Lazarus *et al.,* 2019^90^. Primers used for construction are listed in Supplementary Table 9. We linearized pTOX5 (Genbank MK972845) with the restriction enzyme SwaI, used PCR to amplify ∼1 kb of the *C. rodentium* genome up and downstream of the respective gene/locus (including 2-3 codons on each end), and assembled these three fragments with the HiFi DNA Assembly Master Mix. This construct was electroporated into MFD*pir*, checked by PCR, and conjugated into a streptomycin-resistant strain of *C. rodentium*. *C. rodentium* transconjugants were selected and purified by plating three times on solid LB containing 200 µg/ml streptomycin, 20 µg/ml chloramphenicol, and 0.2% glucose. Single colonies were expanded in liquid LB without selection for 1 h and counter selection was performed on solid LB containing 200 µg/ml streptomycin and 2% rhamnose. The identity of the mutant strains was confirmed by selective plating, PCR, and whole-genome sequencing.

### *C. rodentium* barcoded competitions

*C. rodentium* strains were barcoded using the pSM1 donor library^28^. pSM1 is a library of ∼70,000 plasmids, each with a unique ∼25 bp barcode integrated into a Tn7 transposon. To create barcoded strains, *C. rodentium* was conjugated with MFD*pir*+pSM1 and MFD*pir*+pJMP1039 and multiple, unique barcoded strains were isolated for each mutant. For competition experiments, the barcoded strains were expanded separately in liquid LB to ensure every strain started at an equal concentration.

For competition on solid medium, 18 strains (5 mutants and wild-type, each represented by 3 barcoded strains) were plated on solid LB containing 200 µg/ml streptomycin and incubated at 37 °C. Every 9-15 h the pooled strains were scraped off the plate, resuspended in PBS, diluted, and replated. The size of the population was determined by serial dilution and a sample was taken to determine the relative frequency of each barcode.

For competition in mice, a pool of 6 strains (5 mutants and wild-type) were intragastrically inoculated into mice. Each mouse was given a separate set of 6 barcodes to differentiate the administered strains from bacteria shed by cohoused animals. The total size of the *C. rodentium* population was monitored by serial dilution and a sample was taken from fecal bacteria expanded on solid LB to determine the relative frequency of each barcode.

### Barcode sequencing

The identity and frequency of barcodes were determined by amplicon sequencing as described in Hullahalli *et. al.,* 2021^91^. Bacteria were boiled to release the DNA and PCR was performed to amplify the barcode region. Samples were sequenced on a NextSeq 1000 and the sequencing data was used to determine the number of reads mapping to each barcode. To determine the fold-change in each strain over time, the frequency of each barcode within the sample was normalized to its frequency in the initial population. STAMP sequencing primers are listed in Supplementary Table 9.

## Supporting information

pTn plasmid

pCre plasmid

## Acknowledgements

We thank members of the Waldor lab for helpful discussions and feedback on the manuscript.

## Software

Data analysis was performed using R, Python, and Excel. Graphics and figures were prepared with BioRender, GraphPad Prism and PowerPoint.

## License Information

This article is subject to HHMI’s Open Access to Publications policy. HHMI lab heads have previously granted a nonexclusive CC BY 4.0 license to the public and a sublicensable license to HHMI in their research articles. Pursuant to those licenses, the author-accepted manuscript of this article can be made freely available under a CC BY 4.0 license immediately upon publication.

