## Supplementary figures for "Inducible transposon mutagenesis for genome-scale forward genetics"

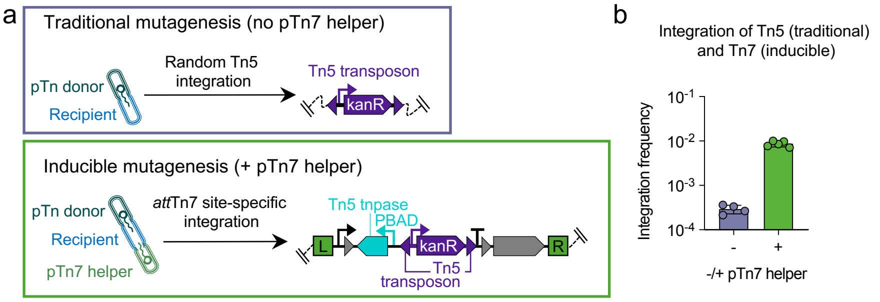


**Extended Data Fig. 1 | Comparison of Tn5 and Tn7 integration frequencies. a**, The non-replicative pTn donor plasmid can be used to create a traditional transposon mutant library through a one-time, random Tn5 transposon integration, mediated by the Tn5 transposase, immediately after the plasmid is introduced into recipient cells. Alternatively, it can be used to create an inducible library by co-introduction of the Tn7 helper plasmid (expressing the Tn7 integration machinery), wherein transposon mutants are generated following the site-specific integration of the Tn5 transposition complex at the *att*Tn7 site in the genome. **b**, Integration at the *att*Tn7 site or random Tn5 integration both result in kanamycin resistance. However, integration is ~30-fold more efficient when the Tn7 helper plasmid is present, indicating that most kanamycin-resistant colonies (total transposon integrants) represent cells containing the Tn5 transposition complex at the *att*Tn7 site rather than random Tn5 mutants. The integration frequency is expressed as the ratio of kanR CFU to total CFU.

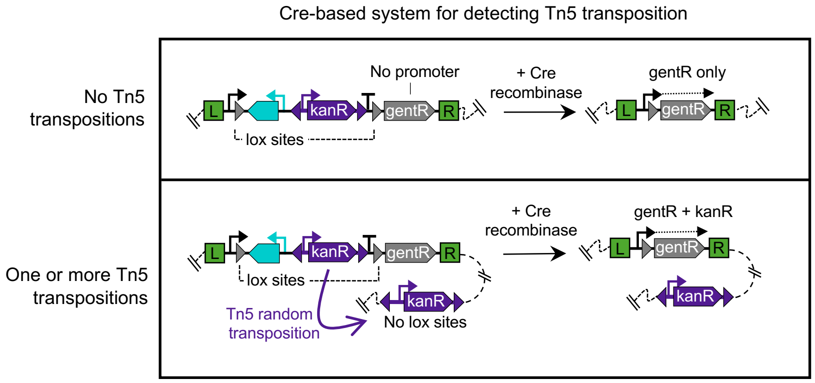


**Extended Data Fig. 2 | A Cre recombinase-based indicator of transposition frequency at the population level.** Following outgrowth of *att*Tn7 site-specific integrants, an optional second conjugation step can be performed to determine the population-level frequency of Tn5 transposition out of the *att*Tn7 site. When a plasmid expressing the Cre recombinase is introduced into the mutant population, Cre expression leads to excision of the Tn5 transposition complex at the attTn7 site via recombination of the lox sequences. Cre excision causes recipient cells to simultaneously lose the attTn7-site kanamycin marker and activate expression of the gentamicin marker. Cells that did not undergo Tn5 transposition prior to Cre excision of the Tn5 transposition complex become solely resistant to gentamicin, while cells that did undergo transposition retain a copy of the Tn5 transposon at a random genomic location outside of the Tn7 lox sites, rendering them resistant to both kanamycin and gentamicin. The ratio of gentR+kanR to gentR colonies provides a measure of transposition frequency in cells where the Tn5 transposition complex was initially integrated at the attTn7 site (Fig. 2b).

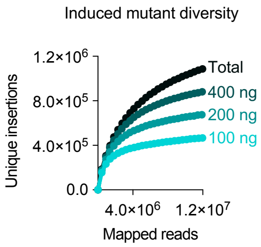


**Extended Data Fig. 3 | Total mutant diversity depends on the amount of DNA sampled.** Increasing the amount of template DNA used in amplification of the InducTn-seq library increases the number of unique insertions detected. The 100 ng sample in this panel is the same as “Induced” in Fig. 2c.

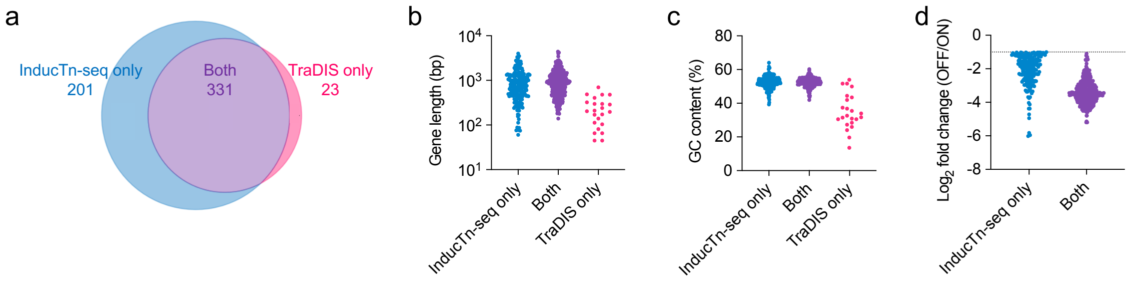


**Extended Data Fig. 4 | Comparison of InducTn-seq to traditional Tn-seq. a,** Venn diagram of *E. coli* MG1655 genes classified as having a fitness defect by InducTn-seq ($\log_2$ fold change <-1 and adjusted *P* value <0.01) and *E. coli* BW25113 genes classified as "essential" by TraDIS. 331 genes were commonly identified between the two screens, representing a 93.5% overlap. The discordant genes identified only by TraDIS were on average (**b**) shorter and (**c**) more AT rich. **d,** Genes exclusively identified by InducTn-seq generally had a weaker fitness defect than genes concordantly identified by the two screens.

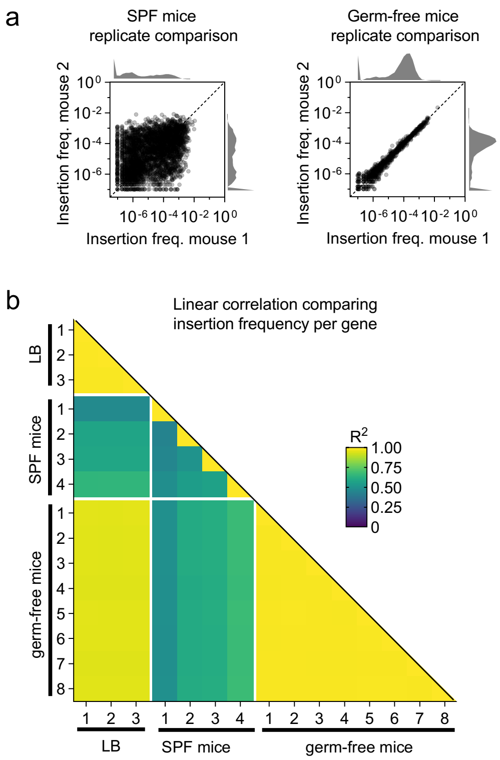


**Extended Data Fig. 5 | Noise in traditional Tn-seq of *C. rodentium* during animal infection is caused by a microbiota-dependent bottleneck.** Tn-seq data from Caballero-Flores et. al. Tn-seq was performed in a traditional manner by administering ~5x$10^4$ unique *C. rodentium* mutants to C57BL/6 mice lacking a microbiota (germ-free) or with an unperturbed, specific pathogen free (SPF) microbiota (bred and maintained at the University of Michigan). Mutants were recovered from the feces of infected animals 2 days post inoculation and sequenced. Consistent with a microbiota-dependent bottleneck randomly removing mutants from the pathogen population, the frequency of *C. rodentium* insertion mutants was more consistent between germ-free than SPF animals. **a**, Points represent genes, insertion frequency represents reads per gene normalized to total reads, and histograms on axes display the distribution of the data. **b**, Coefficient of determination ($R^2$) was used to compare the $\log_{10}$ insertion frequencies.
